# Analyses of circRNA expression throughout circadian rhythm reveal a strong link between Cdr1as and light-induced phase shifts in the SCN

**DOI:** 10.1101/2022.05.18.492346

**Authors:** Andranik Ivanov, Daniele Mattei, Kathrin Radscheit, Anne-Claire Compagnion, J. Patrick Pett, Hanspeter Herzel, Rosa Chiara Paolicelli, Monika Piwecka, Urs Meyer, Dieter Beule

## Abstract

*Cdr1as* is a conserved circular RNA (circRNA) enriched in the CNS and important for maintaining brain homeostasis. The loss of *Cdr1as* results in aberrant synaptic transmission and deregulation of stress response and circadian clock genes. However, it is not known whether the expression of *Cdr1as* or circRNAs, in general, follows a circadian pattern in different tissues. Here, using newly generated and public RNA-Seq data, we monitor circRNA expression throughout circadian rhythm in various mouse brain regions. We demonstrate that *Cdr1as*, despite its stable character, has a highly dynamic expression during the circadian cycle in the mouse suprachiasmatic nucleus (SCN). *Cdr1as* is one of the highest expressed RNAs in a cluster associated with light-induced synaptic transmission and phase shift in the SCN. Further, we identified that another brain enriched circRNA, *mbl*, is also substantially deregulated upon light induction in the fly head. Our study highlights the potential impact of abundant and conserved circRNAs on maintaining a healthy circadian cycle across species.

## Introduction

Circadian rhythms are physiological, molecular, and behavioral changes that follow a 24-hour cycle. Most multicellular organisms take advantage of environmental factors such as light and temperature to fine-tune their daily behavior and metabolism to create a coherent circadian system. In mammals, the suprachiasmatic nucleus (SCN) is the central circadian pacemaker [Hatinsgs *et al*., 2018; Colwell 2015] that is entrained in the light-dark cycles [Schmal *et al*., 2020]. From the retina, the light signal is transmitted to the SCN, which then synchronizes clocks in other tissues [Berson *et al*., 2002]. The endogenous mammalian circadian clock consists of transcription-translation feedback loops [Takahashi, 2017] that are accompanied by global changes in transcriptional and post-transcriptional regulation [Menet *et al*., 2012; Pembroke *et al*., 2015], including RNA synthesis and degradation, its alternative splicing and editing [Terajima *et al*., 2017; Terajima *et al*., 2018; Preussner *et al*., 2014, McGlincy *et al*., 2012]. As of now, it is clear that many RNA species, both protein-coding and non-coding, play an essential role in the maintenance of the circadian rhythm [Menet *et al*., 2012; Pembroke *et al*., 2015; Zhou *et al*., 2021; Kadener *et al*., 2009]. However, circRNAs, an alternative form of RNA splicing products, dependent on temperature and RNA-editing [Ivanov *et al*., 2015; Ashwal-Fluss *et al*., 2014; Rybak-Wolf *et al*., 2015; Shen *et al*., 2022], have not been well studied throughout daily activities and circadian rhythm.

CircRNAs are a large class of highly stable RNA molecules [Salzmann *et al*., 2012; Jeck *et al*., 2013; Memczak *et al*., 2013]. They have conserved biogenesis and are expressed in many animals and plants [Salzman *et al*., 2012; Memczak *et al*., 2013; Ivanov *et al*., 2015; Zhang *et al*., 2014; Ashwal-Fluss *et al*., 2014]. CircRNAs have various functions such as sequestering of miRNAs, modulation of RNA stability, interaction with RNA binding proteins, and regulation of mRNA transcription [Memczak *et al*., 2013; Hansen *et al*., 2013; Ashwal-Fluss *et al*., 2014; Li *et al*., 2015; Du *et al*., 2016]. Certain circRNAs can be translated into proteins [Pamudurti *et al*., 2017; Legnini *et al*., 2017]. In animals, circRNAs are highly expressed in the brain, particularly at the synaptic terminals [Rybak-Wolf *et al*., 2015; You *et al*., 2015]. Due to their high level of stability and synaptic enrichment, these molecules are thought to be involved in intracellular information transport, long-term memory formation, and memory consolidation [Hanan *et al*., 2017]. Moreover, the deregulation of circRNAs has been implicated in neurodegenerative, psychiatric, and neurodevelopmental disorders [Piwecka *et al*., 2017; Zimmerman *et al*., 2020; Zhang *et al*., 2020; Dube *et al*., 2019; Yang *et al*., 2019; Zhang & Brian 2021]. Many of these brain conditions occur in parallel or arguably due to aberrant wake/sleep cycle [Menet & Rosbash 2011; Nasan & Videnovic 2021], which is why it is necessary to study circRNAs in the context of the circadian rhythm.

Here, we studied circRNA expression in different mouse brain regions during 12:12 hour light-dark (LD) cycles and identified *Cdr1as* as an essential regulatory molecule that impacts the light-dependent phase shift in the SCN. *Cdr1as* is highly expressed in the brain, particularly in glutamatergic neurons [Piwecka *et al*., 2017]. It has an unusually high number of miR-7 binding sites [Memczak *et al*., 2013; Hansen *et al*., 2013], a potent, brain-enriched miRNA [Landgraf *et al*., 2007; Zhao *et al*., 2020] that is critical in the pathogenesis of different brain diseases [McMillan *et al*., 2017; Titze-de-Almeida *et al*., 2018; Choi *et al*., 2018; Puthiyedth *et al*., 2016; Madadi *et al*., 2019; Dong *et al*., 2016; Zhao *et al*., 2015; Choi *et al*., 2015; Junn *et al*., 2009]. *Cdr1as* and miR-7 regulate neuronal activity through a network with two other non-coding RNAs, miR-671 and *Cyrano* [Kleaveland *et al*., 2018]. The loss of *Cdr1as* in mice results in deregulation of excitatory synaptic transmission and upregulation of early response and circadian clock genes [Piwecka *et al*., 2017]. *Cdr1as* deletion in mice also leads to vision defects [Chen *et al*., 2020]. Our findings explain previously observed gene expression changes and phenotypes in *Cdr1as* mutant mice and add another piece to the puzzle of *Cdr1as* functions. We further examined overall circRNA expression changes during the LD cycle in fly heads and identified light and period-dependent expression of yet another very conserved and highly abundant circRNA *mbl* [Ashwal-Fluss *et al*., 2014]. Our study opens a new perspective on circRNA functions in circadian rhythm.

## Results

### *Cdr1as* is highly expressed in the SCN and has a significant differential expression during the 12:12h light-dark cycle

We quantified circRNA expression levels in the SCN by systematically analyzing total RNA-Seq data from Pembroke *et al*. [eLife 2015]. The authors monitored six equidistant time points in 9 replicates during a 12:12 hour LD cycle. We identified a total of 1691 expressed circular RNAs. The number of consistently detectable circRNAs was highest in the middle of the dark phase, with a substantial drop at its end (325, 485, 455, 450, 523, and 290 different circRNAs at ZT2, ZT6, ZT10, ZT14, ZT18, and ZT22, respectively, cf. Fig.1A). *Cdr1as* circRNA showed the highest expression of all (Fig. 1B, Supplementary Table 1). It was expressed order of magnitude more than any other circRNA in the SCN (Fig. 1B). *Cdr1as* contributed most to overall differences in circular reads during the whole 12:12 LD cycle (Fig. 1C). It was the only highly expressed circRNA with significant deregulation in SCN over the LD cycle (e.g., ZT14 vs. ZT10: DESeq2 adj.P-value 1.3 e^-19^). A rather unexpected observation is that a stable circRNA is downregulated more than twofold in 4 hours (Fig. 1D) and again upregulated, exhibiting a twin-peak expression with peaks at the beginning and the end of the dark cycle (Fig 1D). Interestingly, Pembroke *et al*. described a cluster of 766 genes with exact twin-peak expression as a synaptic module regulating light-induced phase shifts in the SCN. Inspecting the *Cdr1* locus in the UCSC genome browser showed all reads mapping to the *Cdr1* locus map within the circRNA boundaries rather than the primary transcript (Fig. 1E). Thus all reads mapping to the *Cdr1* gene could be attributed to the circular RNA, allowing a better *Cdr1as* expression quantification (Suppl. Fig. 1A) and putting *Cdr1as* into the top 5 highest (sorted by FPKM) expressed RNAs in the synaptic twin-peak module described by Pembroke *et al*. (Fig. 1F). Variability between replicates differs substantially between time points and is highest near the light-dark transitions, which might hint at quick regulatory processes. Interestingly the long non-coding RNA *Cyrano*, which is known to limit the expression of miR-7 in the brain and thus enables the accumulation of *Cdr1as* [Kleaveland *et al*., 2018], showed the same twin-peak expression pattern as *Cdr1as* (Suppl. Fig. 1, Fig. 1F).

**Figure 1.**
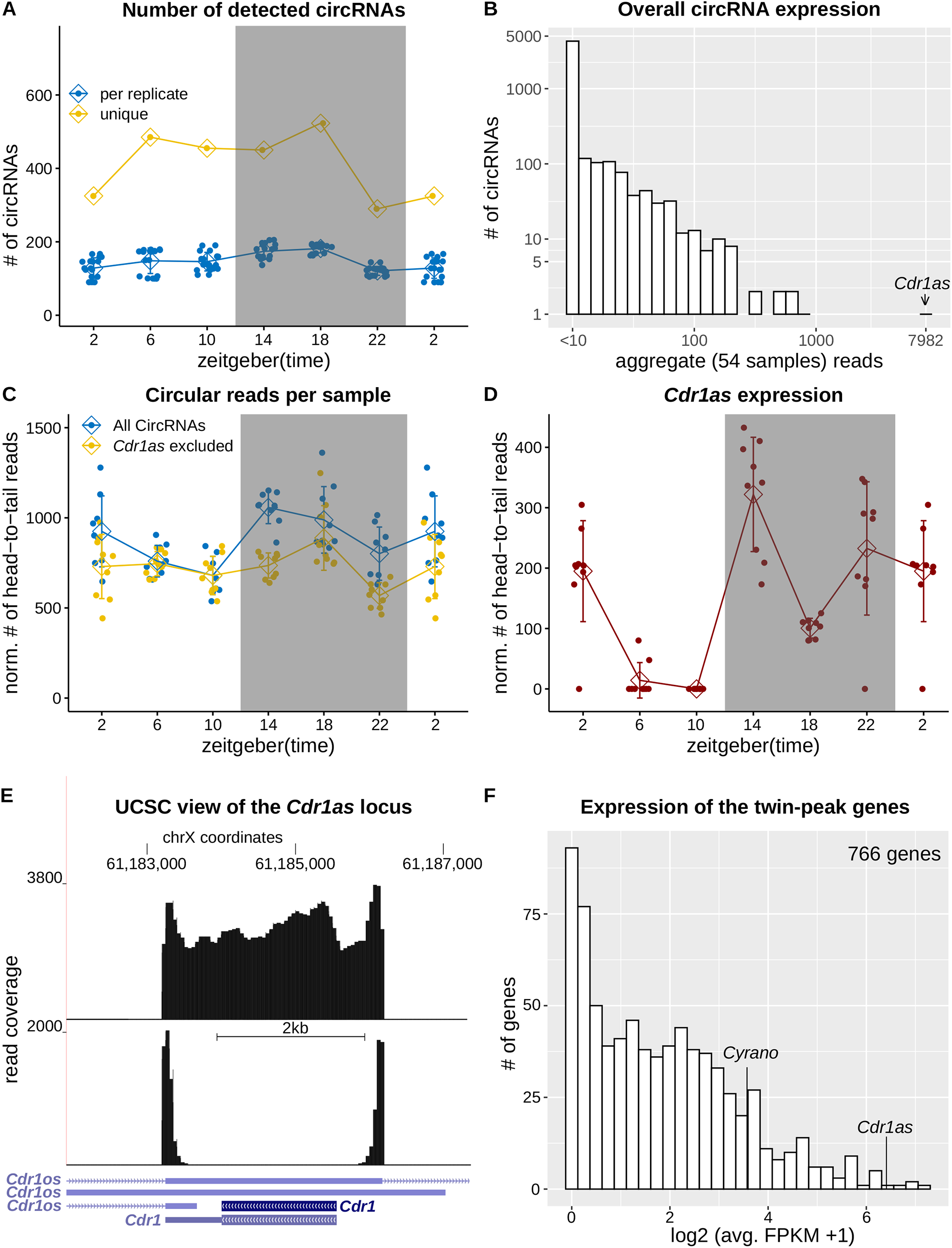
*Cdr1as* expression changes during 24h LD cycle in the SCN. **A)** The number of detected circRNAs for each replicate and each time point. **B)** Overall expression of circRNAs. For each circRNA we summed up circularization supporting head-to-tail reads in all samples/groups. *Cdr1as* has an order of magnitude higher expression than any other circRNA. **C)** Normalized (to the number of mapped reads) circular reads in 12:12 LD cycle. **D)** Normalized expression of *Cdr1as* circRNA during the 12:12 LD cycle. Figure 1A to D use periodic continuation plotting at ZT2. **E)** UCSC genome browser view of the *Cdr1* locus using a pool of reads from three replicates at ZT14. Upper panel shows only circular (head-to-tail) reads. Lower panel shows all reads mapping to the locus. **F)** Histogram shows the expression (FPKM) of all 766 twin-peak module genes from Pembroke *et al*., 2015.

### The expression of *Cdr1as*-associated genes significantly depends on light induction

A study by Piwecka *et al*., 2017 has shown that loss of *Cdr1as* results in upregulation of central clock genes (*Per1, Sik1, Klf10*) and upregulation of immediate early genes such as *Fos, Egr1, Klf4*, Jun, etc. This group of immediate early genes is activated and rapidly transcribed in the SCN upon photic stimuli [Porterfield *et al*., 2007]. Recently, the Takahashi lab measured genome-wide mRNA changes in the SCN after a short period of light exposure (e.g., 30 minutes) [Xu *et al*., 2021]. This early light-induced gene set is significantly enriched in genes deregulated upon *Cdr1as* KO in different brain regions (Fig.2 A, B). We note that despite RNA changes were not measured specifically in the SCNs of *Cdr1as* KO mice, overall gene expression changes are very similar in all investigated brain regions after *Cdr1as* deletion [Piwecka et al., 2017]. Further, Xu *et al*. identified that the *Npas4* transcription factor is an essential regulator of circadian behavior and transcriptional response to light in the SCN [Xu et al., 2021]. In the polyA+ RNA-Seq of the *Cdr1as* KO mice [Piwecka *et al*., 2017], *NPas4* is significantly upregulated in three (hippocampus, cerebellum, olfactory bulb) out of the four sequenced brain regions. Moreover, in the hippocampus (of *Cdr1as* KO mice), *Npas4* is the 11^th^ and the cerebellum’s 7^th^ most significantly deregulated gene.

**Figure 2.**
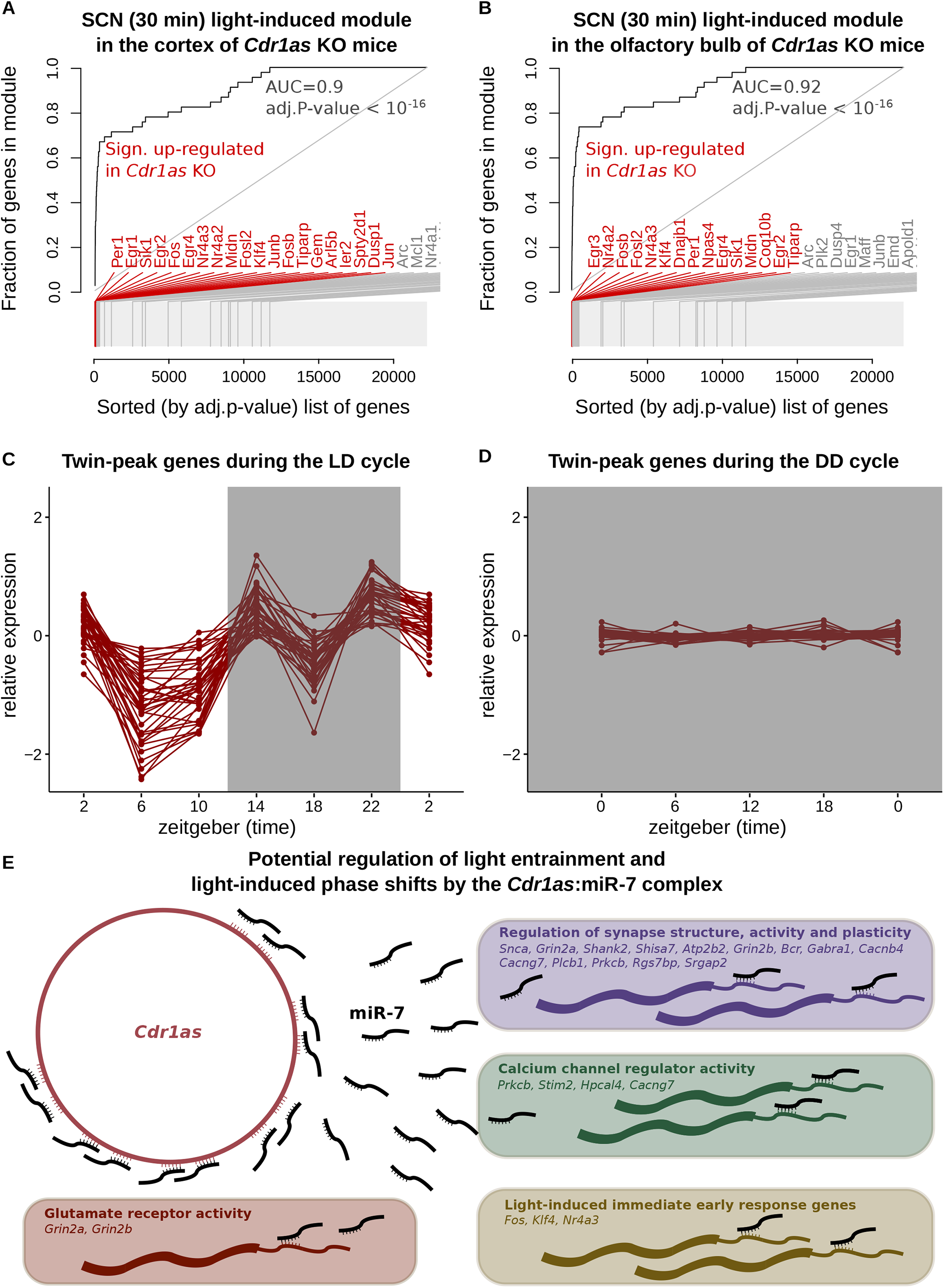
*Cdr1as*/twin-peak regulation and light entrainment. Evidence plots show enrichment of SCN immediate light-induced genes (defined in Xu *et al*., 2021) in **(A)** cortex and **(B)** olfactory bulb of *Cdr1as* KO mice. X-axis shows the list of sorted genes (by adj. p-value *Cdr1as* KO vs WT) from polyA RNA-Seq in Piwecka *et al*., 2017. Y-axis is the fraction of gene-set defined by Xu et al., 2021. In bright red are genes upregulated in *Cdr1as* KO with adj.P-value < 0.05 **C)** Log2 expression (normalized to the temporal mean) of top 50 genes (sorted by FPKM) of the twin-peak module in the LD cycle [Pemborke *et al*., 2015]. **D)** Same set of genes in the DD cycle [Cheng *et al*., 2019]. Cheng *et al*. measured RNA expression in control and Sox2 mutant mice in the DD cycle at four different time points. Here we used only control samples. **E)** *Cdr1as* and miR-7 are both expressed in the SCN. Predicted targets of miR-7 were downloaded from Targetscan [Lewis *et al*., 2005]. Twin-peak genes are defined in Pembroke *et al*.. Immediate light-induced gene-module is from Xu *et al*.. The annotation of other pathway-modules is from MsigDB database.

To further study the effect of light on *Cdr1as* regulation, we analyzed the expression of the twin-peak cluster genes (defined by Pembroke *et al*.) in the SCN during constant darkness using data from [Cheng *et al*., 2019]. The study provides RNAseq at four circadian time points (CT=0,6,12,18) in murine SCN with five replicates each. Before harvesting, the mice were released from the standard 12:12 LD cycle into complete darkness for two days (dark-dark or DD cycle). We selected the top 50 highest expressed genes from the twin-peak module and monitored their expression in LD and DD cycles. Due to the polyA enrichment protocol used by Cheng *et al*., *Cdr1as* circRNA itself is not detectable in this data. However, all other 49 highly expressed genes did not change their expression significantly during the DD cycle (Figure 2C, D). Our observations hold when the analysis is carried out using the top 100 or top 200 highest expressed genes (sorted by FPKM) from the twin-peak module (Suppl. Figure 2). This result suggests that the overall expression of the twin-peak cluster and *Cdr1as* in murine SCN is highly dependent on light exposure.

Light is communicated to the SCN from the retina by glutamatergic neurotransmission from the retinohypothalamic tract. Given the *Cdr1as* enrichment specifically in glutamatergic neurons [Piwecka *et al*., 2017] and high expression of miR-7 in the SCN [Herzer *et al*., 2012; Lee *et al*., 2006], we suggest that *Cdr1as* may be an important regulator of the glutamatergic synaptic transmission during the light-induced phase-shifts through the delivery of miR-7 to its early response and twin-peak target genes. The overlap of miR-7 targets (444 in total from the Targetscan database [Lewis *et al*., 2005]) and twin-peak cluster (766) identified 37 genes (Suppl. Table 2) that could be regulated by *Cdr1as*/miR-7 complex in the context of light-induced phase-shifts. Pembroke *et al*. reported that the twin-peak cluster is enriched in synaptic transmission and calcium signaling. As expected, its subset miR-7 targets are also significantly enriched in these biological pathways (Suppl. Table 2). Here we infer the *Cdr1as*:miR-7 complex’s role in regulating photic inputs in the SCN (Fig. 2E). *Cdr1as* may regulate miR-7 target genes from the twin-peak cluster that form ionotropic glutamate receptor complex: Grin2a and Grin2b. *Cdr1as*:miR-7 can also impact calcium influx (via *Prkcb, Stim2, Hpcal4*, and *Cacng7*) and subsequent generation of nitric oxide, resulting in different responses depending on the time of the day [Pembroke *et al*., 2015]. Finally, *Cdr1as*/miR-7 complex may regulate the synaptic structure and maintain its activity through *Snca, Grin2a, Shank2, Shisa7, Atp2b2, Grin2b, Bcr, Gabra1* (Figure 2E).

### CircRNA expression in the hippocampus and frontal cortex follows constitutive transcription patterns

To further characterize the circadian expression of circRNAs, we sequenced total RNA from two additional brain regions, the hippocampus and frontal cortex. While the SCN acts as a circadian master pacemaker, other brain regions also display oscillatory capacity [Chun *et al*., 2015]. The hippocampus is particularly interesting as it is essential for sleep-dependent memory consolidation [Squire *et al*., 1991]. It is interesting to study circRNA regulation in this context as circRNAs are highly expressed in the hippocampus [Rybak-Wolf *et al*., 2015] and have a much longer lifetime compared to linear RNAs, and therefore, may be involved in memory consolidation as well as transport of information between different cell types [Hanan *et al*., 2017]. We also investigated the frontal cortex as it is associated with many psychiatric and neurodegenerative disorders.

Using three replicates at six-time points (SE 150bp, average sequencing depth 46 mil.), we identified 5505 and 4790 distinct circRNAs expressed in the hippocampus and frontal lobe (Suppl. Table 3). Our analysis in the frontal cortex identified 1328 of 1770 (75%) *circbase* circRNAs [*circbase*: Glazar *et al*., 2014], and in the hippocampus, 1297 of 1676 circbase circRNAs. In the hippocampus, overall circRNA expression decreased by 20 percent during the transition from the dark to the light phase (Fig 3B). However, such a change in circularized RNA was most probably due to the difference in the expression of the linear host transcripts (Fig. 3A, Suppl. Fig. 3). The expression of *Cdr1as* in the hippocampus was downregulated by 30% during the transition from the light to the dark phase and again upregulated during the dark phase. In the frontal cortex, *Cdr1as* expression did not change significantly during the LD cycle (Fig. 3C). The overall circRNA expression in the cortex increased by 30 percent in the middle of the wake phase and was downregulated again just before the light phase started. Similar to the hippocampus, such a fluctuation of circularization was due to a general change of the read fraction falling in exonic regions (Fig. 3B, Suppl. Fig. 3).

**Figure 3.**
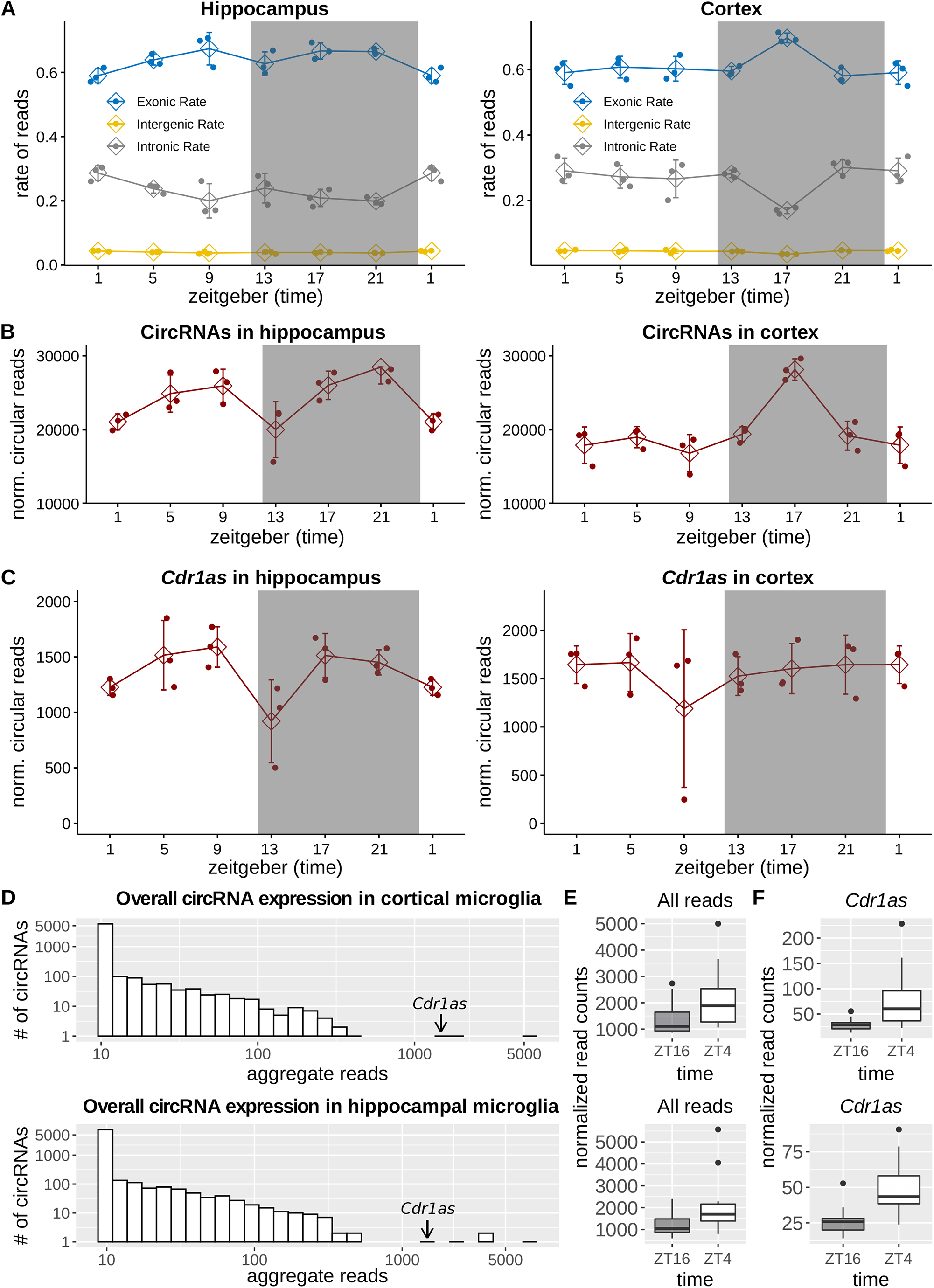
CircRNAs in the hippocampus and frontal cortex. **A)** Genomic annotation of the sequencing reads. The X-axis is the (zeitgeber) time and Y-axis is the fraction of reads. B) Normalized (norm. factor = total number of reads/avg. library size) number of circular reads per time point. **C)** Normalized (norm. factor = total number of reads/avg. library size) number of circular reads supporting *Cdr1as*. **D)** Analogous to Fig. 1B. **E)** Normalized (norm. factor = total number of reads/avg. library size) number of all circular reads detected in hippocampal (down) and cortical (up) microglia sequencing. Each group represents a distribution of 16 replicates. **F)** *Cdr1as* expression (normalized circular reads) in microglia from cortex (upper panel) and hippocampus (lower panel). Each box 16 samples.

### *Cdr1as* is the only significantly deregulated circRNA between day and night in microglia

Microglia are brain resident macrophages, and their inflammatory responses are controlled by the intrinsic circadian clock [Fonken *et al*., 2015]. Microglia also exhibit morphological differences between wake and sleep, and disruption of the clock system of these cells may result in impaired behavior and contribute to the disturbance of sleep [Nakanishi *et al*., 2021]. The regulation of circRNAs in microglia is of particular interest as these cells are in close contact with synapses and, like neuronal cells, express genes with long introns that predominantly give rise to circRNAs [Jeck *et al*., 2013; Ivanov *et al*., 2015]. Besides producing circRNAs, microglia could phagocytose circRNAs that are exported by neurons to the synaptic space. We sequenced freshly isolated microglia from the mouse hippocampus and frontal cortex during the light and dark phases: ZT4 and ZT16 (per time-point 16 replicates, SE 150bp, average sequencing depth: 42 mil.). Our analyses identified 1234 and 1553 distinct circRNAs in the cortex and hippocampus, respectively (Suppl. Table 4). *Cdr1as* was one of the highest expressed circRNAs in microglia (Fig 3D). Despite an overall increase in circularization during the light phase (Fig. 3E), the only circRNA that significantly changed its expression in both cortex and hippocampus was *Cdr1as* (Fig 3F). In Cortex, *Cdr1as* upregulated during the light phase by ∼2.6 fold (DESeq2: adj. P-value=1.6*10^−7^) and in the hippocampus by 1.8 fold (DESeq2: adj. P-value=1.6*10^−7^).

### Analysis of circRNAs in the fly head reveals light-dependent expression of circ-*mbl*

To further study the circadian regulation of circRNAs across species, we analyzed their expression in the fly brain over the 12:12 LD cycle. Earlier research described the splicing patterns and expression of non-coding RNAs in the fly brain during the LD cycle [Hughes *et al*., 2012]. The study has monitored the transcriptome changes along the 12:12 LD cycle in wild-type and *period*-null clock-defective animals. Alternative splicing throughout time did not change significantly in fly heads. However, over 600 significantly alternatively spliced exons were detected in *per*-null animals [Hughes *et al*., 2012]. CircRNAs are mostly non-coding splicing isoforms highly expressed in neuronal cells and progressively accumulate during aging [Rybak-Wolf *et al*., 2015; You *et al*., 2015; Westholm *et al*., 2014]. Thus we analyzed their expression in the Hughes et al. data set. Consistently with alternative splicing results, the expression of circRNAs did not change much over time. From highly expressed circRNAs only circ-mbl substantially (∼3 fold) changed its expression after light induction (Fig. 4B), the second most highly expressed circRNA in the fly brain (Fig. 4A). *Mbl* circRNA is one of the most studied circRNAs expressed in humans, mice, and flies [Ashwal-Fluss *et al*., 2014]. Besides its high degree of conservation [Ashwal-Fluss *et al*., 2014], it is of particular interest as this is one of the few circRNAs that have shown the ability to give rise to protein [Pamudurti *et al*., 2017]. Circ-*mbl* is a multifunctional molecule, and its downregulation leads to male developmental lethality, altered gene expression, and behavioral defects [Pamudurti *et al*., bioarchive 2018].

**Figure 4.**
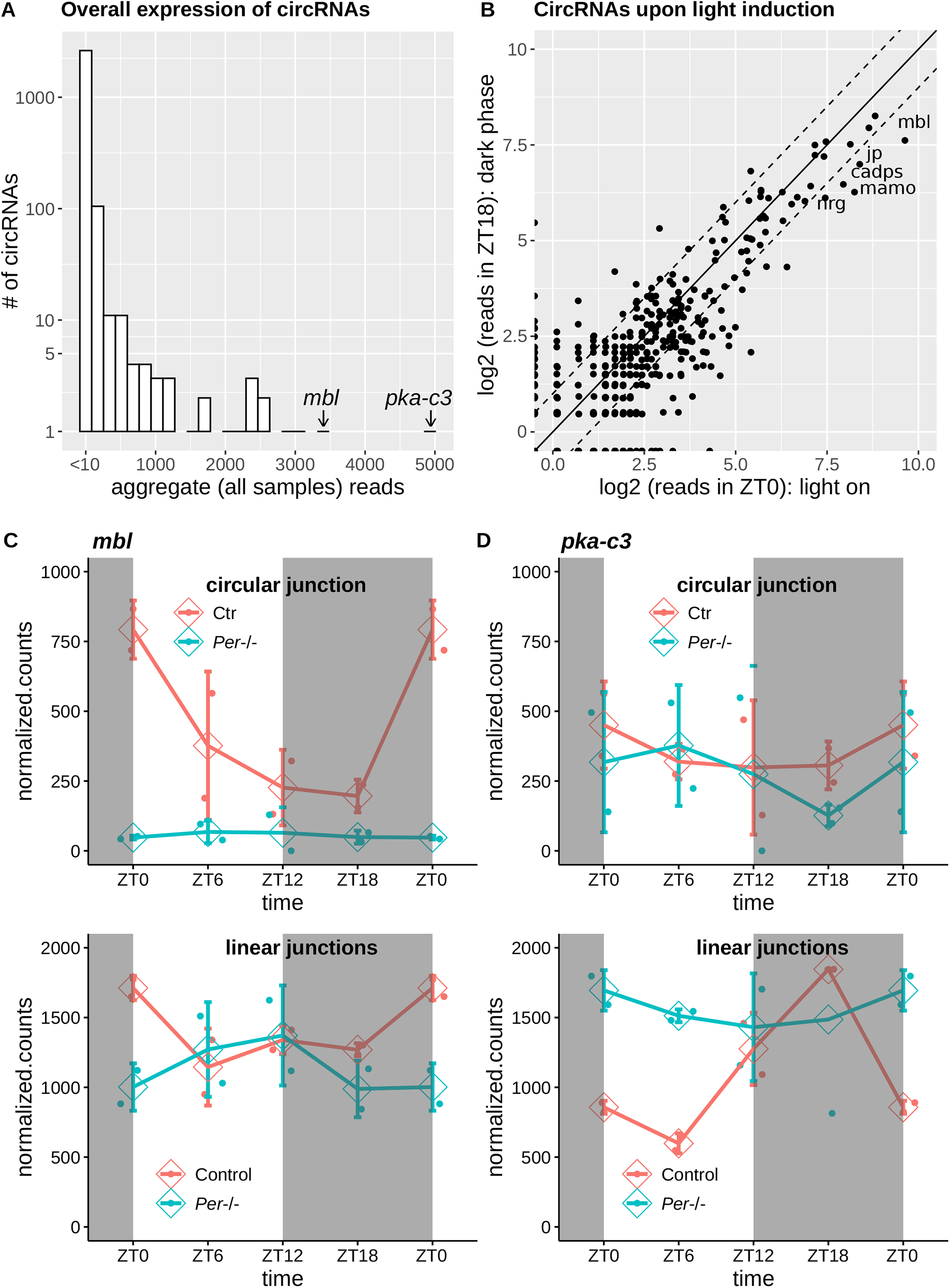
CircRNA expression throughout circadian rhythm in fly. CircRNA expression in fly heads throughout the circadian rhythm. **A)** Analogous to Fig 1B. **B)** Comparison of circRNA expression in ZT18 (dark) and ZT0 (light on) time points. The expression values are normalized to the total number of reads. Shown are the average values over 2 replicates. **C)** Normalized expression of circRNAs (up) and linear (down) RNA in control (red) and *per-*mutant (green) mice. **D)** Data for *pka-c3*, analogous to figure 5C.

Circ-*mbl* expression decreased more than threefold from the beginning of the light cycle to the end of the dark cycle. However, the expression of the linear isoform decreased only 1.5 fold in this period (Fig. 4C). In *Per-*deficient flies, the expression of circ-*mbl* almost disappeared while the linear isoform did not change its expression (Fig. 4C). Circ-*pka-c3* was found to have the highest expression in the fly head. This circRNA has shown an opposite behavior to circ-mbl. While its linear isoform increased its expression threefold from the light to dark cycle, the circular isoform did not change (Fig. 4D). This result suggests that circRNAs may be regulated in a circadian manner independent of their host RNA.

## Discussion

Our study identifies two very abundant and conserved circular RNAs that have light and perhaps activity-dependent expression patterns in animals. We characterize *Cdr1as* circRNA as a novel gene associated with light-entrainment in the SCN. This rationalizes why in all four investigated brain regions (cortex, hippocampus, cerebellum, and olfactory bulb), Piwecka *et al*. recorded significant upregulation of core clock genes in *Cdr1as* deficient mice. What was unknown at the time of this study is that loss of *Cdr1as* also caused downregulation of a miRNA family (miR-96/-182/-183) that was recently discovered to be an important modulator of circadian rhythm [Zhou *et al*., 2021]. Although there is no evidence that photic impulse directly regulates *Cdr1as* expression, we show that genes upregulated upon *Cdr1as* deletion are highly enriched in the light-induced immediate response pathway. Moreover, we demonstrate that the strong regulation of the synaptic twin-peak cluster during the 12 hours of the dark cycle (wake phase for mice) occurs only after withdrawal from the light cycle. In the retina, deletion of *Cdr1as* circRNA results in increased beta-wave amplitude of the photopic electrophysiological response and reduced vision contrast sensitivity [Chen *et al*., 2020]. Together these observations suggest that *Cdr1as* plays an important role in communicating light from the retina to SCN, most likely through the regulation of glutamatergic neurotransmission.

Since two other members of the regulatory network described by Kleaveland *et al*., miR-7 and *Cyrano* are also expressed in the SCN, the central circadian pacemaker emerges as a perfect system to study the interplay of these non-coding molecules. One surprising observation in the SCN is that *Cdr1as*, a highly stable circular RNA, is downregulated by more than 2 fold during 4 hours (from ZT14 to ZT18). One possible mechanism may be the cleavage of circRNA by miR-671 [Hansen *et al*., 2011]. Also, *Cdr1as* turnover may occur as a result of structure-mediated RNA decay by *Upf1* and *G3bp1* [Fischer *et al*., 2020]. Both genes are highly expressed in the SCN.

Pembroke *et al*. reported that the twin-peak module is highly enriched with genes involved in synaptic transmission, long-term potentiation, calcium signaling, and gated channel activity. On the other hand, Piwecka et al. demonstrated that the loss of *Cdr1as* circRNA in mice results in aberrant excitatory synaptic transmission. We believe that further experiments exploring the exact pathway by which *Cdr1as* dampens neuronal function should investigate the presence of *Cdr1as* with miR-7 targets from the twin-peak module (Figure 2, Supplementary Table 2). Particularly interesting is the interaction of *Cdr1as*/miR-7 with alpha-synuclein (Snca) (twin-peak gene). Abnormal expression of *Snca* and its aggregation is critical in the pathophysiology of Parkinson’s disease (PD) [Stefanis 2012]. Moreover, in PD patients, miR-7 is significantly downregulated in the brain regions that undergo dopaminergic neurodegeneration during the course of the disease [McMillan *et al*., 2017]. Due to its binding ability to *Snca* 3’UTR in vivo and subsequent regulation of protein translation, miR-7 is becoming widely appreciated as an important therapeutic target for PD [Junn *et al*., 2009; Choi *et al*., 2018; Titze-de-Almeida *et al*., 2018; McMillan *et al*., 2017]. Sleep disorders are a common feature of patients in the early stages of the disease. Thus, understanding the strong regulation of *Cdr1as* in the central circadian pacemaker may provide a new paradigm for studying the early onset of PD. It is important to highlight that the present study demonstrates oscillations in circRNA expression patterns in physiological conditions in adult animals. As research is developing around circRNA dynamics in brain development and specific disorders, it will be highly relevant to screen how the latter are affected in disease from a circadian perspective. In preclinical studies, mice are most often sacrificed for downstream molecular analyses during their sleep phase, while in clinical studies, the time of collection of samples from human patients and controls is rarely taken into consideration as a biologically relevant variable. Hence, when studying transcriptional dynamics of both coding and non-coding RNAs, implementing a circadian component will widen our understanding of transcriptional (de)regulation. This concept might be helpful, especially when studying *Cdr1as* in psychiatric conditions and neurodegenerative diseases, where altered sleep and circadian patterns are a disease hallmark [Freeman *et al*., 2020; Fifel and Videnovic, 2020].

## Methods

### Animals

Nine to ten weeks old male C57BL6/N mice (Charles River Laboratories, Sulzfeld, Germany) were used throughout the study. We allocated 3-5 animals per cage in individually ventilated cages (IVCs). The animal vivarium was a specific-pathogen-free (SPF) holding room, which was temperature- and humidity-controlled (21 ± 3 °C, 50 ± 10%) Animals used for the microglial isolation were kept under a reversed light-dark cycle (lights off: 09:00 AM–09.00 PM), whilst the animals used for whole hippocampal and frontal lobe tissue sequencing were kept on normal light-dark cycle (Lights on: 6 AM-6 PM). All animals had *ad libitum* access to the same food (Kliba 3436, Kaiseraugst, Switzerland) and water throughout the entire study. All procedures described in the present study had been previously approved by the Cantonal Veterinarian’s Office of Zurich, and all efforts were made to minimize the number of animals used and their suffering.

### Brain Dissociation and Cell Isolation

Brain tissue dissociation and microglia cell isolation were performed according to a protocol recently optimized and published by us [Mattei & Ivanov *et al*., 2021]. The protocol is carried out at 4C to avoid cell activation due to the isolation procedure. Briefly, the animals were deeply anesthetized with an overdose of Nembutal (Abbott Laboratories, North Chicago, IL, USA) and transcardially perfused with 15 ml ice-cold, calcium- and magnesium-free Dulbecco’s phosphate-buffered saline (DPBS, pH 7.3-7.4). The brains were quickly removed and washed with ice-cold DPBS, after which the hippocampi and frontal cortices were dissected on a cooled petri dish and placed in ice-cold Hibernate-A medium. Mechanical dissociation (MD) at 4°C was carried out on ice. The tissue was dissociated in 1.5 ml Hibernate-A medium in a 1 ml Dounce homogenizer with a loose pestle. The homogenized tissue was then sieved through a 70μm cell strainer. The homogenates were pelleted at 450xg for 6 minutes at 4°C. The supernatants were removed, and the pellets were re-suspended with a P1000 micropipette, applying a pipette-tip cut-off. 500 μL of freshly prepared isotonic percoll solution was then added to each sample (final volume: 2 ml) and mixed well. Percoll was rendered isotonic by mixing 1 part of 10x calcium- and magnesium-free DPBS (pH 7.3-7.4) with 9-parts of percoll. Importantly, the pH of percoll was adjusted to 7.3-7.4 with 5 molar hydrochloric acid before starting the isolation procedure. The percoll solution was mixed properly with the cell suspension, after which 2 ml of DPBS were gently layered on top of it with a pipette boy set on the slowest speed, creating two separate layers. The samples were centrifuged for 10 minutes at 3000xg. The centrifugation resulted in an upper layer consisting of DPBS and a lower layer consisting of percoll. The two layers were separated by a disk of myelin and debris, while the cells were located at the bottom of the tube. The layers were aspirated, leaving about 500 μL. The cells were then washed once in DPBS and pelleted by centrifuging them at 460xg for10 minutes at 4°C. This pellet consists of total brain cells, including microglial cells.

### Microglia Cell Magnetic Sorting

Microglia cells were isolated via magnetic-activated cell sorting (MACS) using mouse anti-CD11b magnetic microbeads (Miltenyi) according to the manufacturer’s instructions with some modifications. The MACS buffer used consisted of 2% bovine serum albumin (BSA) diluted in DPBS from a 7.5% cell-culture grade BSA stock (Thermo Fisher Scientific). Total hippocampal/frontal cortex cell pellets after percoll separation (see above) were re-suspended in 90 μL MACS buffer and 10 μL anti-mouse-CD11b magnetic beads (Miltenyi). The cells were then incubated for 15 min at 4°C. Cells were washed with 1 ml MACS buffer and pelleted at 300 rcf for 5 min at 4°C. The cells were then passed through an MS MACS column (Miltenyi) attached to a magnet. After washing the columns three times with MACS buffer, microglia were flushed from the column with 1 ml MACS buffer and pelleted at 300 rcf for 5 min at 4°C. Cell pellets were then snap-frozen in liquid nitrogen and stored at −80°C.

### Hippocampal and Frontal Lobe Tissue Collection

For the total RNA-seq of whole brain tissue, RNA was extracted from adult male mice total hippocampi and frontal lobes. Briefly, deeply anesthetized adult male mice were intracardially perfused with ice-cold Dulbecco’s phosphate-buffered saline (DPBS) to remove blood. Hippocampi and frontal lobes were dissected immediately after in a pre-cooled sterile Petri dish on ice. The brain regions were immediately transferred to an RNAse-free Eppendorf tube, snap-frozen in liquid nitrogen, and stored at −80 °C until RNA extraction.

### Total RNA Extraction

Total RNA from freshly isolated microglia and from hippocampal and frontal lobe tissue was extracted via phenol/chloroform extraction using the SPLIT-RNA extraction kit (Lexogen, product code: 008.48) according to the manufacturer’s instructions. The RNA was treated with Turbo DNase I (Ambion, product code AM1907) to remove traces of genomic DNA. Following DNase I treatment, the RNA was stored at −80 °C until library preparation.

### Total RNA Library Preparation and Sequencing

Before library preparation, the integrity of each sample was assessed on an Agilent TapeStation system 4150 using the RNA screen tape (Agilent). In contrast, RNA concentrations were measured on a Qubit 4 fluorometer (ThermoFisher Scientific). 100 ng total RNA were used as input for ribosomal RNA (rRNA) depletion using the NEBNext rRNA depletion kit (New England BioLabs inc., product code: E6350) according to the manufacturer’s instructions. Following rRNA depletion, total RNA libraries were built using the NEBNext Ultra II library prep kit for Illumina (New England BioLabs inc., product code: E7775) according to the manufacturer’s instructions. The yield of amplified libraries was measured on a Qubit 4 fluorometer using the Qubit high sensitivity DNA kit (HS DNA kit). Amplified libraries were further analyzed on the HS D1000 screen tape on a TapeStation system 4150 to assess library size and molarity prior to pooling. The libraries were sequenced using Illumina HISeq-4000.

### Bioinformatic methods

For circRNA detection, total RNA seq reads were mapped to the mouse GRCm38 or drosophila dm6 genomes with BWA [Li & Durbin 2009] (version 0.7.17-r1188) using the -T 19 option. CircRNA were identified using CIRI2 [Gao *et al*., 2015] with default parameters. For further analyses, we used circRNAs identified in at least two replicates of the same time point. To compare circRNAs with circbase datasets, circbase coordinates were translated from mm9 to mm10 genome using the UCSC liftOver tool. For the calculation of the p-values and LFCs for circRNA expression changes, we used DESeq2 [Love *et al*., 2014] (version 1.22.1) package (normalization: rld, test: Wald’s test, p-value adjustment: Benjamini-Hochberg). The circRNA expression table was concatenated with the linear constitutive gene expression table. To calculate the expression of the linear genes, we mapped the reads to the mouse GRCm38 genome with STAR (version 2.7.3.a, default parameters, [Dobin *et al*., 2013]) and assigned reads to genes with featureCounts [Liao *et al*., 2014] (version 2.0.0). For the FPKM calculation, read counts were normalized to the total number of uniquely mapped reads per sample and the length of the genes as reported by featureCounts (calculated from the gencode GRCm38 vM12 GTF file). For the comparison of the linear and circular RNA expression changes in the hippocampus and cortex, we quantified the sum of all spliced reads mapped to the gene (featureCounts -J parameter). The evidence plots for Figure 2A, B were produced with R tmod (version 0.46.2 [Zyla *et al*., 2019]). Gene ontology analysis for Supplementary Table 2 and Figure 2E were carried out with R tmod and msigdbr (7.4.1) packages.

## Author Contributions

Conceptualization, A.I., D.M.; Bioinformatic Analyses, A.I.; Experiments D.M., A.C.C., and M.P.; Funding Acquisition U.M., D.B., D.M.; Supervision, A.I., R.C.P., U.M., and D.B.; Writing - Original Draft, A.I.; Visualization A.I.; Writing - Review & Editing, A.I., D.M., K.R., P.P., M.P., H.H., R.C.P., and D.B.; Investigation A.I., D.M., K.R., A.C.C., P.P., M.P., and D.B., Project Administration A.I., and D.B

## Acknowledgments

We thank Benedikt Obermayer from BIH Core Unit Bioinformatics for insightful comments on the manuscript. The authors also thank Tatiana Borodina from the MDC/BIH genomics facility for the library preparation and RNASeq.

## Declaration of interests

The authors declare no competing interests.

## Code and data availability

The raw sequencing data have been deposited in GEO under accession numbers GSE199791 and GSE200314 and will be available upon publication. The R code for data analyses and figures can be downloaded from github.com/bihealth/circ_circ

## Figure legends

**Supplementary Figure 1.**
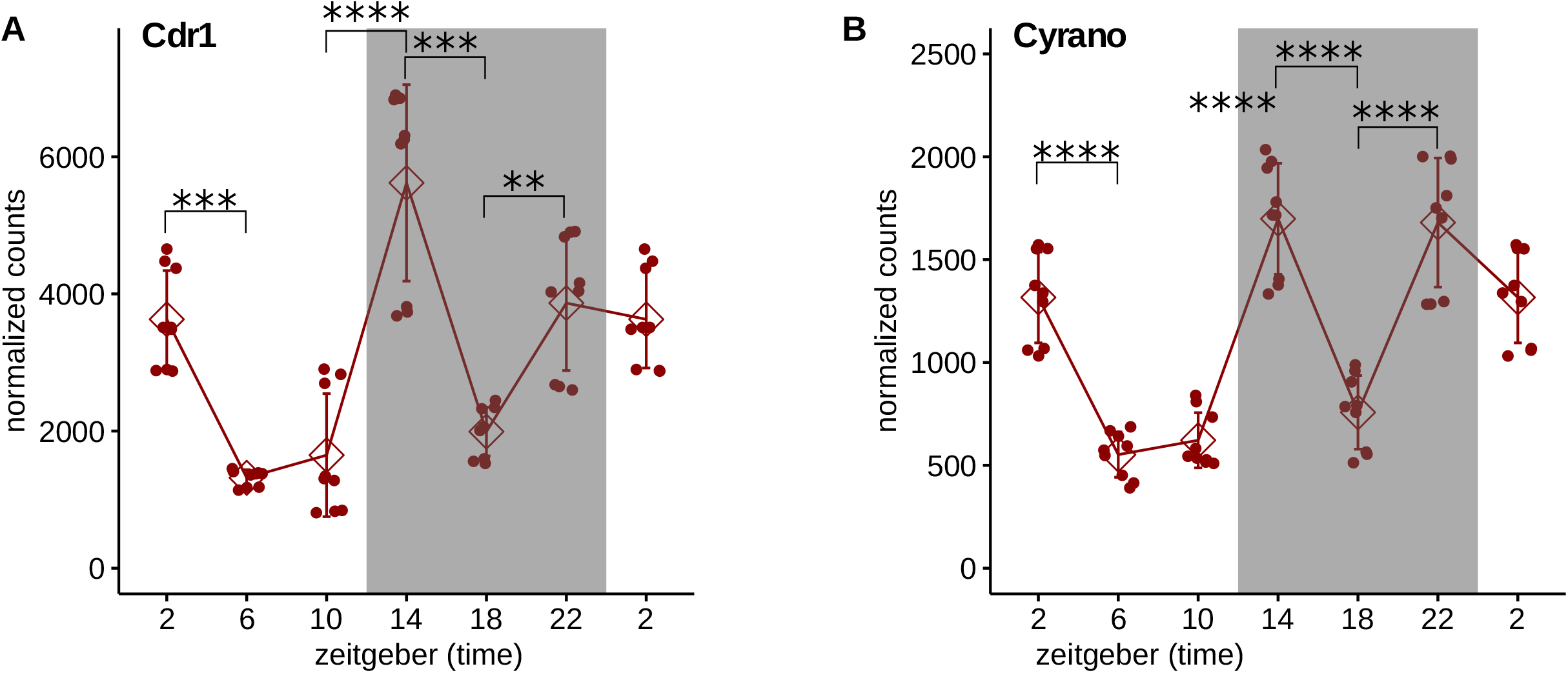
*Cdr1as* and *Cyrano* expression in the SCN. Normalized (norm. factor = total number of reads/avg. library size) read counts of *Cdr1as* (including linear reads) and *Cyrano* non-coding RNA. *Cdr1* coordinates in the GTF file were modified to the start and end coordinates of the *Cdr1as* circRNA. DESeq2 Benjamini-Hochberg adjusted p-values: ** < 10^−3^, *** < 10^−6,^ **** < 10^−10^.

**Supplementary Figure 2.**
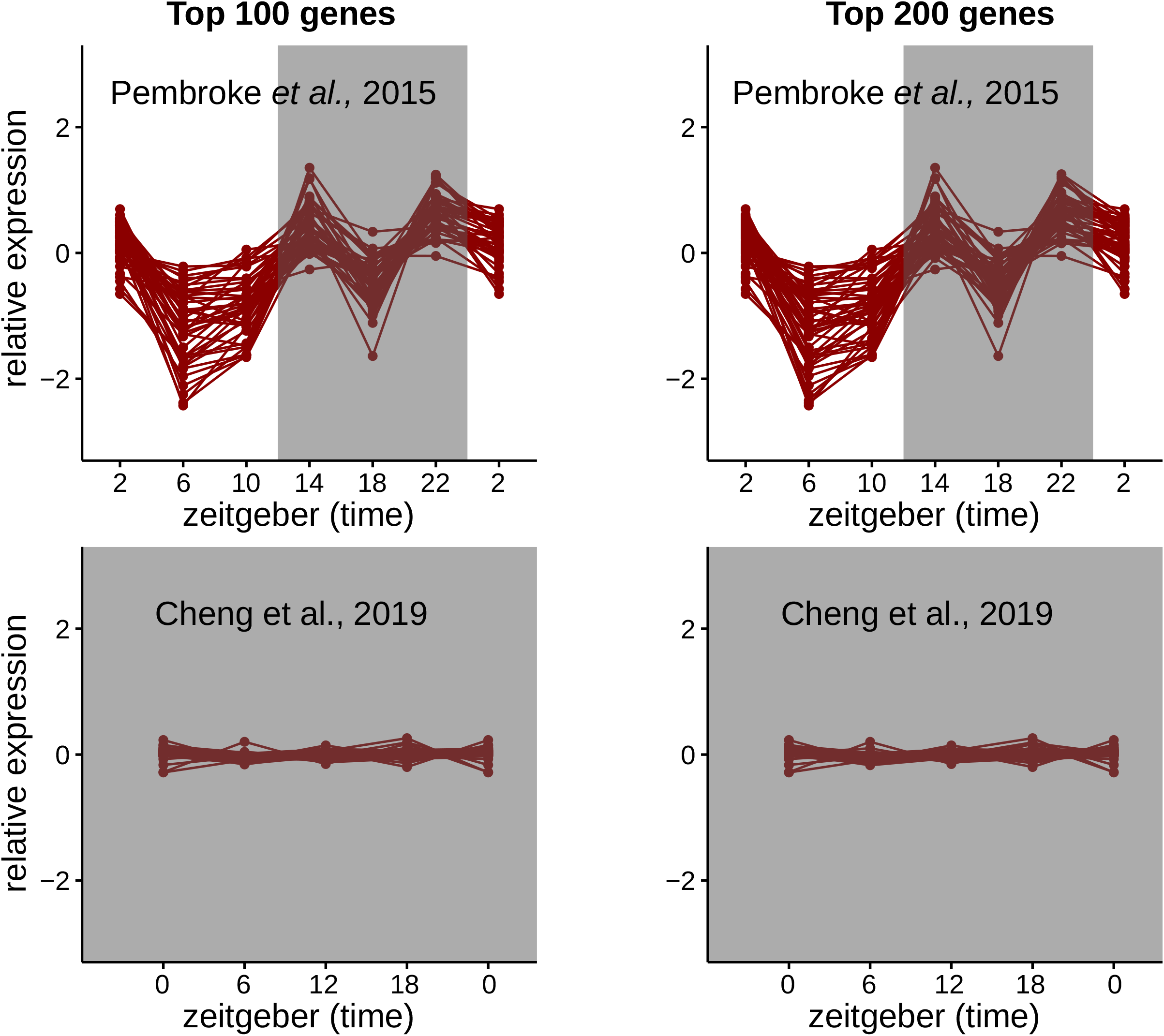
Twin peak cluster during the DD and LD cycles. top 100 and top 200 highly expressed genes from the twin peak module: analogous to Figure 2 C,D

**Supplementary Figure 3.**
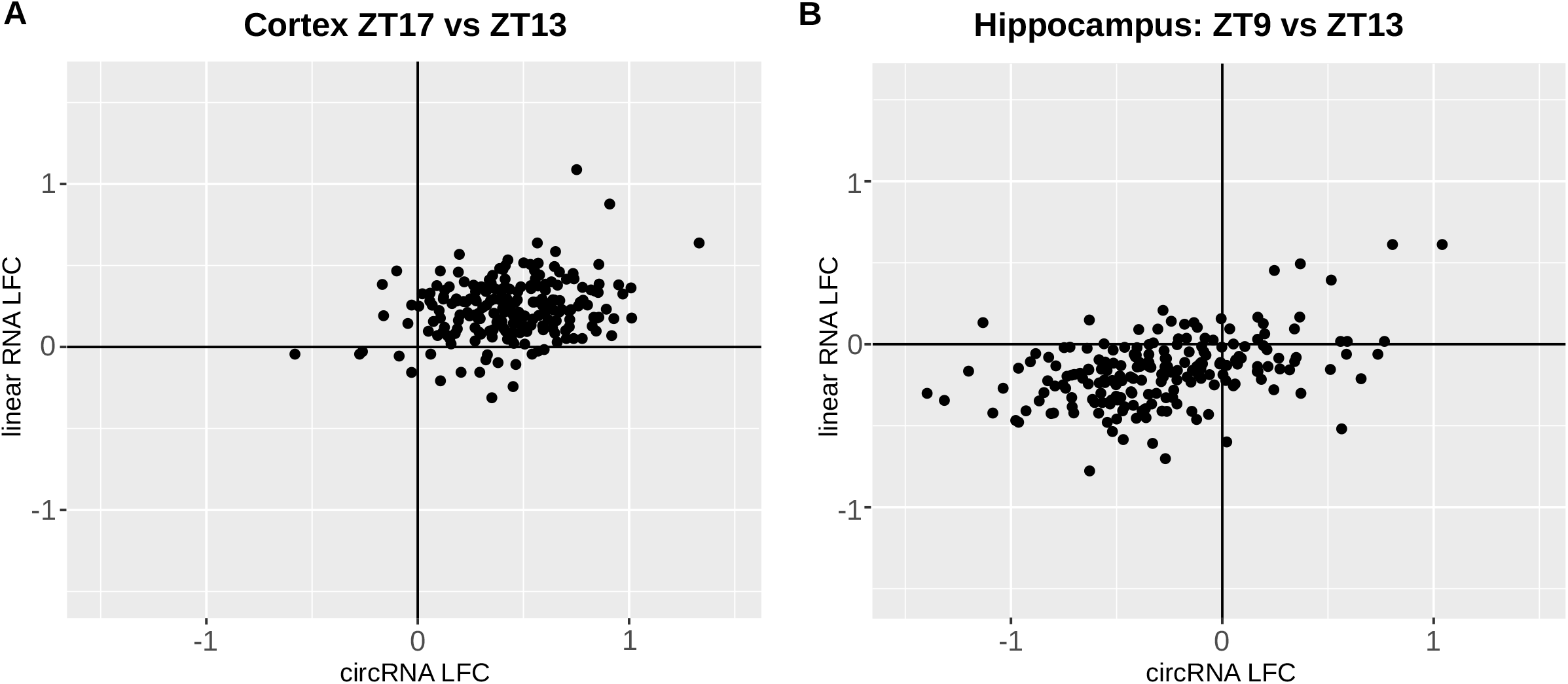
Dependence of CircRNA expression on their linear isoforms. Comparison of circRNA expression changes with expression changes of their linear host transcripts. X-axis is the log2 fold change of circRNAs (pseudocount=5). Y-axis is the log fold change of the circRNA host linear isoform (sum of linear reads mapping to the splice sites). We count only reads mapping to the linear splice junctions of the gene. Pseudocount=5. We considered only circRNAs that have on average more than 30 reads in 2 time points: In cortex avg.ZT17+avg.ZT13>30 and in the hippocampus avg.ZT9+avg.ZT13>30.

## Notes

### Competing Interest Statement

The authors have declared no competing interest.

## References

Ashwal-Fluss, R., Meyer, M., Pamudurti, N. R., Ivanov, A., Bartok, O., Hanan, M., Evantal, N., Memczak, S., Rajewsky, N., & Kadener, S. (2014). circRNA biogenesis competes with pre-mRNA splicing. Molecular Cell, 56(1), 55–66. https://doi.org/10.1016/j.molcel.2014.08.019

Berson, D. M., Dunn, F. A., & Takao, M. (2002). Phototransduction by retinal ganglion cells that set the circadian clock. Science (New York, N.Y.), 295(5557), 1070–1073. https://doi.org/10.1126/science.1067262

Chen, X.-J., Li, M.-L., Wang, Y.-H., Mou, H., Wu, Z., Bao, S., Xu, Z.-H., Zhang, H., Wang, X.-Y., Zhang, C.-J., Xue, X., & Jin, Z.-B. (2020). Abundant Neural circRNA Cdr1as Is Not Indispensable for Retina Maintenance. Frontiers in Cell and Developmental Biology, 8, 565543. https://doi.org/10.3389/fcell.2020.565543

Cheng, A. H., Bouchard-Cannon, P., Ness, R. W., & Cheng, H.-Y. M. (2019). RNA-sequencing data highlighting the time-of-day-dependent transcriptome of the central circadian pacemaker in Sox2-deficient mice. Data in Brief, 24, 103909. https://doi.org/10.1016/j.dib.2019.103909

Choi, D. C., Yoo, M., Kabaria, S., & Junn, E. (2018). MicroRNA-7 facilitates the degradation of alpha-synuclein and its aggregates by promoting autophagy. Neuroscience Letters, 678, 118–123. https://doi.org/10.1016/j.neulet.2018.05.009

Choi, S.-Y., Pang, K., Kim, J. Y., Ryu, J. R., Kang, H., Liu, Z., Kim, W.-K., Sun, W., Kim, H., & Han, K. (2015). Post-transcriptional regulation of SHANK3 expression by microRNAs related to multiple neuropsychiatric disorders. Molecular Brain, 8(1), 74. https://doi.org/10.1186/s13041-015-0165-3

Chun, L. E., Woodruff, E. R., Morton, S., Hinds, L. R., & Spencer, R. L. (2015). Variations in Phase and Amplitude of Rhythmic Clock Gene Expression across Prefrontal Cortex, Hippocampus, Amygdala, and Hypothalamic Paraventricular and Suprachiasmatic Nuclei of Male and Female Rats. Journal of Biological Rhythms, 30(5), 417–436. https://doi.org/10.1177/0748730415598608

Colwell, C. S. (2011). Linking neural activity and molecular oscillations in the SCN. Nature Reviews. Neuroscience, 12(10), 553–569. https://doi.org/10.1038/nrn3086

Dong, Y.-F., Chen, Z.-Z., Zhao, Z., Yang, D.-D., Yan, H., Ji, J., & Sun, X.-L. (2016). Potential role of microRNA-7 in the anti-neuroinflammation effects of nicorandil in astrocytes induced by oxygen-glucose deprivation. Journal of Neuroinflammation, 13(1), 60. https://doi.org/10.1186/s12974-016-0527-5

Dobin, A., Davis, C. A., Schlesinger, F., Drenkow, J., Zaleski, C., Jha, S., Batut, P., Chaisson, M., & Gingeras, T. R. (2013). STAR: ultrafast universal RNA-seq aligner. Bioinformatics (Oxford, England), 29(1), 15–21. https://doi.org/10.1093/bioinformatics/bts635

Du, W. W., Yang, W., Liu, E., Yang, Z., Dhaliwal, P., & Yang, B. B. (2016). Foxo3 circular RNA retards cell cycle progression via forming ternary complexes with p21 and CDK2. Nucleic Acids Research, 44(6), 2846–2858. https://doi.org/10.1093/nar/gkw027

Dube, U., Del-Aguila, J. L., Li, Z., Budde, J. P., Jiang, S., Hsu, S., Ibanez, L., Fernandez, M. V., Farias, F., Norton, J., Gentsch, J., Wang, F., Dominantly Inherited Alzheimer Network (DIAN), Salloway, S., Masters, C. L., Lee, J.-H., Graff-Radford, N. R., Chhatwal, J. P., Bateman, R. J., … Cruchaga, C. (2019). An atlas of cortical circular RNA expression in Alzheimer disease brains demonstrates clinical and pathological associations. Nature Neuroscience, 22(11), 1903–1912. https://doi.org/10.1038/s41593-019-0501-5

Fifel, K., & Videnovic, A. (2020). Circadian and Sleep Dysfunctions in Neurodegenerative Disorders-An Update. Frontiers in Neuroscience, 14, 627330. https://doi.org/10.3389/fnins.2020.627330

Fischer, J. W., Busa, V. F., Shao, Y., & Leung, A. K. L. (2020). Structure-Mediated RNA Decay by UPF1 and G3BP1. Molecular Cell, 78(1), 70-84.e6. https://doi.org/10.1016/j.molcel.2020.01.021

Fonken, L. K., Frank, M. G., Kitt, M. M., Barrientos, R. M., Watkins, L. R., & Maier, S. F. (2015). Microglia inflammatory responses are controlled by an intrinsic circadian clock. Brain, Behavior, and Immunity, 45, 171–179. https://doi.org/10.1016/j.bbi.2014.11.009

Freeman, D., Sheaves, B., Waite, F., Harvey, A. G., & Harrison, P. J. (2020). Sleep disturbance and psychiatric disorders. The Lancet. Psychiatry, 7(7), 628–637. https://doi.org/10.1016/S2215-0366(20)30136-X

Gao, Y., Wang, J., & Zhao, F. (2015). CIRI: an efficient and unbiased algorithm for de novo circular RNA identification. Genome Biology, 16, 4. https://doi.org/10.1186/s13059-014-0571-3

Glažar, P., Papavasileiou, P., & Rajewsky, N. (2014). circBase: a database for circular RNAs. RNA (New York, N.Y.), 20(11), 1666–1670. https://doi.org/10.1261/rna.043687.113

Ivanov, A., Memczak, S., Wyler, E., Torti, F., Porath, H. T., Orejuela, M. R., Piechotta, M., Levanon, E. Y., Landthaler, M., Dieterich, C., & Rajewsky, N. (2015). Analysis of intron sequences reveals hallmarks of circular RNA biogenesis in animals. Cell Reports, 10(2), 170–177. https://doi.org/10.1016/j.celrep.2014.12.019

Hanan, M., Soreq, H., & Kadener, S. (2017). CircRNAs in the brain. RNA Biology, 14(8), 1028–1034. https://doi.org/10.1080/15476286.2016.1255398

Hastings, M. H., Maywood, E. S., & Brancaccio, M. (2018). Generation of circadian rhythms in the suprachiasmatic nucleus. Nature Reviews. Neuroscience, 19(8), 453–469. https://doi.org/10.1038/s41583-018-0026-z

Hansen, T. B., Jensen, T. I., Clausen, B. H., Bramsen, J. B., Finsen, B., Damgaard, C. K., & Kjems, J. (2013). Natural RNA circles function as efficient microRNA sponges. Nature, 495(7441), 384–388. https://doi.org/10.1038/nature11993

Hansen, T. B., Wiklund, E. D., Bramsen, J. B., Villadsen, S. B., Statham, A. L., Clark, S. J., & Kjems, J. (2011). miRNA-dependent gene silencing involving Ago2-mediated cleavage of a circular antisense RNA. The EMBO Journal, 30(21), 4414–4422. https://doi.org/10.1038/emboj.2011.359

Herzer, S., Silahtaroglu, A., & Meister, B. (2012). Locked Nucleic Acid-Based In Situ Hybridisation Reveals miR-7a as a Hypothalamus-Enriched MicroRNA with a Distinct Expression Pattern. Journal of Neuroendocrinology, 24(12), 1492–1504. https://doi.org/10.1111/j.1365-2826.2012.02358.x

Hughes, M. E., Grant, G. R., Paquin, C., Qian, J., & Nitabach, M. N. (2012). Deep sequencing the circadian and diurnal transcriptome of Drosophila brain. Genome Research, 22(7), 1266–1281. https://doi.org/10.1101/gr.128876.111

Jeck, W. R., Sorrentino, J. A., Wang, K., Slevin, M. K., Burd, C. E., Liu, J., Marzluff, W. F., & Sharpless, N. E. (2013). Circular RNAs are abundant, conserved, and associated with ALU repeats. RNA (New York, N.Y.), 19(2), 141–157. https://doi.org/10.1261/rna.035667.112

Junn, E., Lee, K.-W., Jeong, B. S., Chan, T. W., Im, J.-Y., & Mouradian, M. M. (2009). Repression of alpha-synuclein expression and toxicity by microRNA-7. Proceedings of the National Academy of Sciences of the United States of America, 106(31), 13052–13057. https://doi.org/10.1073/pnas.0906277106

Kadener, S., Menet, J. S., Sugino, K., Horwich, M. D., Weissbein, U., Nawathean, P., Vagin, V. v, Zamore, P. D., Nelson, S. B., & Rosbash, M. (2009). A role for microRNAs in the Drosophila circadian clock. Genes & Development, 23(18), 2179–2191. https://doi.org/10.1101/gad.1819509

Kleaveland, B., Shi, C. Y., Stefano, J., & Bartel, D. P. (2018). A Network of Noncoding Regulatory RNAs Acts in the Mammalian Brain. Cell, 174(2), 350-362.e17. https://doi.org/10.1016/j.cell.2018.05.022

Landgraf, P., Rusu, M., Sheridan, R., Sewer, A., Iovino, N., Aravin, A., Pfeffer, S., Rice, A., Kamphorst, A. O., Landthaler, M., Lin, C., Socci, N. D., Hermida, L., Fulci, V., Chiaretti, S., Foà, R., Schliwka, J., Fuchs, U., Novosel, A., … Tuschl, T. (2007). A mammalian microRNA expression atlas based on small RNA library sequencing. Cell, 129(7), 1401–1414. https://doi.org/10.1016/j.cell.2007.04.040

Lee, H.-J., Palkovits, M., & Young, W. S. (2006). miR-7b, a microRNA up-regulated in the hypothalamus after chronic hyperosmolar stimulation, inhibits Fos translation. Proceedings of the National Academy of Sciences of the United States of America, 103(42), 15669–15674. https://doi.org/10.1073/pnas.0605781103

Legnini, I., di Timoteo, G., Rossi, F., Morlando, M., Briganti, F., Sthandier, O., Fatica, A., Santini, T., Andronache, A., Wade, M., Laneve, P., Rajewsky, N., & Bozzoni, I. (2017). Circ-ZNF609 Is a Circular RNA that Can Be Translated and Functions in Myogenesis. Molecular Cell, 66(1), 22- 37.e9. https://doi.org/10.1016/j.molcel.2017.02.017

Lewis, B. P., Burge, C. B., & Bartel, D. P. (2005). Conserved seed pairing, often flanked by adenosines, indicates that thousands of human genes are microRNA targets. Cell, 120(1), 15–20. https://doi.org/10.1016/j.cell.2004.12.035

Li, H., & Durbin, R. (2009). Fast and accurate short read alignment with Burrows-Wheeler transform. Bioinformatics (Oxford, England), 25(14), 1754–1760. https://doi.org/10.1093/bioinformatics/btp324

Li, Z., Huang, C., Bao, C., Chen, L., Lin, M., Wang, X., Zhong, G., Yu, B., Hu, W., Dai, L., Zhu, P., Chang, Z., Wu, Q., Zhao, Y., Jia, Y., Xu, P., Liu, H., & Shan, G. (2015). Exon-intron circular RNAs regulate transcription in the nucleus. Nature Structural & Molecular Biology, 22(3), 256–264. https://doi.org/10.1038/nsmb.2959

Liao, Y., Smyth, G. K., & Shi, W. (2014). featureCounts: an efficient general purpose program for assigning sequence reads to genomic features. Bioinformatics (Oxford, England), 30(7), 923–930. https://doi.org/10.1093/bioinformatics/btt656

Love, M. I., Huber, W., & Anders, S. (2014). Moderated estimation of fold change and dispersion for RNA-seq data with DESeq2. Genome Biology, 15(12), 550. https://doi.org/10.1186/s13059-014-0550-8

Madadi, S., Schwarzenbach, H., Saidijam, M., Mahjub, R., & Soleimani, M. (2019). Potential microRNA-related targets in clearance pathways of amyloid-β: novel therapeutic approach for the treatment of Alzheimer’s disease. Cell & Bioscience, 9(1), 91. https://doi.org/10.1186/s13578-019-0354-3

Mattei, D., Ivanov, A., van Oostrum, M., Pantelyushin, S., Richetto, J., Mueller, F., Beffinger, M., Schellhammer, L., vom Berg, J., Wollscheid, B., Beule, D., Paolicelli, R. C., & Meyer, U. (2020). Enzymatic Dissociation Induces Transcriptional and Proteotype Bias in Brain Cell Populations. International Journal of Molecular Sciences, 21(21), 7944. https://doi.org/10.3390/ijms21217944

McGlincy, N. J., Valomon, A., Chesham, J. E., Maywood, E. S., Hastings, M. H., & Ule, J. (2012). Regulation of alternative splicing by the circadian clock and food related cues. Genome Biology, 13(6), R54. https://doi.org/10.1186/gb-2012-13-6-r54

McMillan, K. J., Murray, T. K., Bengoa-Vergniory, N., Cordero-Llana, O., Cooper, J., Buckley, A., Wade-Martins, R., Uney, J. B., O’Neill, M. J., Wong, L. F., & Caldwell, M. A. (2017). Loss of MicroRNA-7 Regulation Leads to α-Synuclein Accumulation and Dopaminergic Neuronal Loss In Vivo. Molecular Therapy, 25(10), 2404–2414. https://doi.org/10.1016/j.ymthe.2017.08.017

Memczak, S., Jens, M., Elefsinioti, A., Torti, F., Krueger, J., Rybak, A., Maier, L., Mackowiak, S. D., Gregersen, L. H., Munschauer, M., Loewer, A., Ziebold, U., Landthaler, M., Kocks, C., le Noble, F., & Rajewsky, N. (2013). Circular RNAs are a large class of animal RNAs with regulatory potency. Nature, 495(7441), 333–338. https://doi.org/10.1038/nature11928

Menet, J. S., & Rosbash, M. (2011). When brain clocks lose track of time: cause or consequence of neuropsychiatric disorders. Current Opinion in Neurobiology, 21(6), 849–857. https://doi.org/10.1016/j.conb.2011.06.008

Menet, J. S., Rodriguez, J., Abruzzi, K. C., & Rosbash, M. (2012). Nascent-Seq reveals novel features of mouse circadian transcriptional regulation. ELife, 1, e00011. https://doi.org/10.7554/eLife.00011

Nakanishi, H., Ni, J., Nonaka, S., & Hayashi, Y. (2021). Microglial circadian clock regulation of microglial structural complexity, dendritic spine density and inflammatory response. Neurochemistry International, 142, 104905. https://doi.org/10.1016/j.neuint.2020.104905

Nassan, M., & Videnovic, A. (2022). Circadian rhythms in neurodegenerative disorders. Nature Reviews. Neurology, 18(1), 7–24. https://doi.org/10.1038/s41582-021-00577-7

Pamudurti, N.R., Konakondla-Jacob, V.V., Krishnamoorthy, A., Ashwal-Fluss, R., Bartok, O., Wuerst, S., Seitz, K., Maya, R., Lerner, N., Patop, I.L., Rizzoli, S., Beatus, T., Kadener, S. (2018). An in vivo knockdown strategy reveals multiple functions for circMbl. Bioarchive https://doi.org/10.1101/483271

Pamudurti, N.R., Bartok, O., Jens, M., Ashwal-Fluss, R., Stottmeister, C., Ruhe, L., Hanan, M., Wyler, E., Perez-Hernandez, D., Ramberger, E., Shenzis, S., Samson, M., Dittmar, G., Landthaler, M., Chekulaeva, M., Rajewsky, N., & Kadener, S. (2017). Translation of CircRNAs. Molecular Cell, 66(1), 9-21.e7. https://doi.org/10.1016/j.molcel.2017.02.021

Pembroke, W. G., Babbs, A., Davies, K. E., Ponting, C. P., & Oliver, P. L. (2015). Temporal transcriptomics suggest that twin-peaking genes reset the clock. ELife, 4. https://doi.org/10.7554/eLife.10518

Piwecka, M., Glažar, P., Hernandez-Miranda, L. R., Memczak, S., Wolf, S. A., Rybak-Wolf, A., Filipchyk, A., Klironomos, F., Cerda Jara, C. A., Fenske, P., Trimbuch, T., Zywitza, V., Plass, M., Schreyer, L., Ayoub, S., Kocks, C., Kühn, R., Rosenmund, C., Birchmeier, C., & Rajewsky, N. (2017). Loss of a mammalian circular RNA locus causes miRNA deregulation and affects brain function. Science (New York, N.Y.), 357(6357). https://doi.org/10.1126/science.aam8526

Porterfield, V. M., Piontkivska, H., & Mintz, E. M. (2007). Identification of novel light-induced genes in the suprachiasmatic nucleus. BMC Neuroscience, 8, 98. https://doi.org/10.1186/1471-2202-8-98

Preußner, M., Wilhelmi, I., Schultz, A.-S., Finkernagel, F., Michel, M., Möröy, T., & Heyd, F. (2014). Rhythmic U2af26 alternative splicing controls PERIOD1 stability and the circadian clock in mice. Molecular Cell, 54(4), 651–662. https://doi.org/10.1016/j.molcel.2014.04.015

Puthiyedth, N., Riveros, C., Berretta, R., & Moscato, P. (2016). Identification of Differentially Expressed Genes through Integrated Study of Alzheimer’s Disease Affected Brain Regions. PLOS ONE, 11(4), e0152342. https://doi.org/10.1371/journal.pone.0152342

Rybak-Wolf, A., Stottmeister, C., Glažar, P., Jens, M., Pino, N., Giusti, S., Hanan, M., Behm, M., Bartok, O., Ashwal-Fluss, R., Herzog, M., Schreyer, L., Papavasileiou, P., Ivanov, A., Öhman, M., Refojo, D., Kadener, S., & Rajewsky, N. (2015). Circular RNAs in the Mammalian Brain Are Highly Abundant, Conserved, and Dynamically Expressed. Molecular Cell, 58(5), 870–885. https://doi.org/10.1016/j.molcel.2015.03.027

Salzman, J., Gawad, C., Wang, P. L., Lacayo, N., & Brown, P. O. (2012). Circular RNAs are the predominant transcript isoform from hundreds of human genes in diverse cell types. PloS One, 7(2), e30733. https://doi.org/10.1371/journal.pone.0030733

Schmal, C., Herzel, H., & Myung, J. (2020). Clocks in the Wild: Entrainment to Natural Light. Frontiers in Physiology, 11, 272. https://doi.org/10.3389/fphys.2020.00272

Shen, H., An, O., Ren, X., Song, Y., Tang, S. J., Ke, X.-Y., Han, J., Tay, D. J. T., Ng, V. H. E., Molias, F. B., Pitcheshwar, P., Leong, K. W., Tan, K.-K., Yang, H., & Chen, L. (2022). ADARs act as potent regulators of circular transcriptome in cancer. Nature Communications, 13(1), 1508. https://doi.org/10.1038/s41467-022-29138-2

Squire, L. R., & Zola-Morgan, S. (1991). The medial temporal lobe memory system. Science (New York, N.Y.), 253(5026), 1380–1386. https://doi.org/10.1126/science.1896849

Stefanis, L. (2012). α-Synuclein in Parkinson’s disease. Cold Spring Harbor Perspectives in Medicine, 2(2), a009399. https://doi.org/10.1101/cshperspect.a009399

Takahashi, J. S. (2017). Transcriptional architecture of the mammalian circadian clock. Nature Reviews. Genetics, 18(3), 164–179. https://doi.org/10.1038/nrg.2016.150

Terajima, H., Yoshitane, H., Ozaki, H., Suzuki, Y., Shimba, S., Kuroda, S., Iwasaki, W., & Fukada, Y. (2017). ADARB1 catalyzes circadian A-to-I editing and regulates RNA rhythm. Nature Genetics, 49(1), 146–151. https://doi.org/10.1038/ng.3731

Terajima, H., Yoshitane, H., Yoshikawa, T., Shigeyoshi, Y., & Fukada, Y. (2018). A-to-I RNA editing enzyme ADAR2 regulates light-induced circadian phase-shift. Scientific Reports, 8(1), 14848. https://doi.org/10.1038/s41598-018-33114-6

Titze-de-Almeida, R., & Titze-de-Almeida, S. S. (2018). miR-7 Replacement Therapy in Parkinson’s Disease. Current Gene Therapy, 18(3), 143–153. https://doi.org/10.2174/1566523218666180430121323

Westholm, J. O., Miura, P., Olson, S., Shenker, S., Joseph, B., Sanfilippo, P., Celniker, S. E., Graveley, B. R., & Lai, E. C. (2014). Genome-wide analysis of drosophila circular RNAs reveals their structural and sequence properties and age-dependent neural accumulation. Cell Reports, 9(5), 1966–1980. https://doi.org/10.1016/j.celrep.2014.10.062

Xu, P., Berto, S., Kulkarni, A., Jeong, B., Joseph, C., Cox, K. H., Greenberg, M. E., Kim, T.-K., Konopka, G., & Takahashi, J. S. (2021). NPAS4 regulates the transcriptional response of the suprachiasmatic nucleus to light and circadian behavior. Neuron, 109(20), 3268-3282.e6. https://doi.org/10.1016/j.neuron.2021.07.026

Yang, L., Yang, F., Zhao, H., Wang, M., & Zhang, Y. (2019). Circular RNA circCHFR Facilitates the Proliferation and Migration of Vascular Smooth Muscle via miR-370/FOXO1/Cyclin D1 Pathway. Molecular Therapy. Nucleic Acids, 16, 434–441. https://doi.org/10.1016/j.omtn.2019.02.028

You, X., Vlatkovic, I., Babic, A., Will, T., Epstein, I., Tushev, G., Akbalik, G., Wang, M., Glock, C., Quedenau, C., Wang, X., Hou, J., Liu, H., Sun, W., Sambandan, S., Chen, T., Schuman, E. M., & Chen, W. (2015). Neural circular RNAs are derived from synaptic genes and regulated by development and plasticity. Nature Neuroscience, 18(4), 603–610. https://doi.org/10.1038/nn.3975

Zhang, X.-O., Wang, H.-B., Zhang, Y., Lu, X., Chen, L.-L., & Yang, L. (2014). Complementary sequence-mediated exon circularization. Cell, 159(1), 134–147. https://doi.org/10.1016/j.cell.2014.09.001

Zhang, Y., Du, L., Bai, Y., Han, B., He, C., Gong, L., Huang, R., Shen, L., Chao, J., Liu, P., Zhang, H., Zhang, H., Gu, L., Li, J., Hu, G., Xie, C., Zhang, Z., & Yao, H. (2020). CircDYM ameliorates depressive-like behavior by targeting miR-9 to regulate microglial activation via HSP90 ubiquitination. Molecular Psychiatry, 25(6), 1175–1190. https://doi.org/10.1038/s41380-018-0285-0

Zhang, M., & Bian, Z. (2021). The Emerging Role of Circular RNAs in Alzheimer’s Disease and Parkinson’s Disease. Frontiers in Aging Neuroscience, 13, 691512. https://doi.org/10.3389/fnagi.2021.691512

Zhao, H., Xu, J., Pang, L., Zhang, Y., Fan, H., Liu, L., Liu, T., Yu, F., Zhang, G., Lan, Y., Bai, J., Li, X., & Xiao, Y. (2015). Genome-wide DNA methylome reveals the dysfunction of intronic microRNAs in major psychosis. BMC Medical Genomics, 8(1), 62. https://doi.org/10.1186/s12920-015-0139-4

Zhao, J., Zhou, Y., Guo, M., Yue, D., Chen, C., Liang, G., & Xu, L. (2020). MicroRNA-7: expression and function in brain physiological and pathological processes. Cell & Bioscience, 10(1), 77. https://doi.org/10.1186/s13578-020-00436-w

Zhou, L., Miller, C., Miraglia, L. J., Romero, A., Mure, L. S., Panda, S., & Kay, S. A. (2021). A genome-wide microRNA screen identifies the microRNA-183/96/182 cluster as a modulator of circadian rhythms. Proceedings of the National Academy of Sciences of the United States of America, 118(1). https://doi.org/10.1073/pnas.2020454118

Zimmerman, A. J., Hafez, A. K., Amoah, S. K., Rodriguez, B. A., Dell’Orco, M., Lozano, E., Hartley, B. J., Alural, B., Lalonde, J., Chander, P., Webster, M. J., Perlis, R. H., Brennand, K. J., Haggarty, S. J., Weick, J., Perrone-Bizzozero, N., Brigman, J. L., & Mellios, N. (2020). A psychiatric disease-related circular RNA controls synaptic gene expression and cognition. Molecular Psychiatry, 25(11), 2712–2727. https://doi.org/10.1038/s41380-020-0653-4

Zyla, J., Marczyk, M., Domaszewska, T., Kaufmann, S. H. E., Polanska, J., & Weiner, J. (2019). Gene set enrichment for reproducible science: comparison of CERNO and eight other algorithms. Bioinformatics (Oxford, England), 35(24), 5146–5154. https://doi.org/10.1093/bioinformatics/btz447

